# Deep learning analyses of DNA sequences resolve the retention of the Duffy-null resistance to Plasmodium vivax malaria in Africa

**DOI:** 10.64898/2025.12.25.695976

**Authors:** Guillaume Laval, Alice Decugis, Oğuzhan Parasayan, Etienne Patin, Lluis Quintana-Murci, Jacques Chiaroni

## Abstract

P. *vivax*, the most geographically widespread human malaria parasite with millions of clinical cases per year, is however quasi absent in sub-Saharan Africa. Positive selection targeting the rs2814778 protective mutation, also known as the Duffy-null allele, may explain the absence (or quasi absence) of vivax in sub-Saharan Africa by a progressive purge of the pathogen due to a quasi-fixation of the Duffy-null allele and the resulting high rates of protected carriers in western, central and eastern populations. Yet, while positive selection has been clearly evidenced in admixed populations coexisting with vivax, the selection model currently admitted poorly explains the lack of the Duffy-null allele in Europe, or in Asia where the pathogen is mainly observed. In this article, several validated Deep Learning methods applied to high coverage sequence data obtained in 589 African individuals resolved this retention of the Duffy-null resistance to vivax in Africa. The CNN and GAN algorithms implemented in this study also predict a rise in frequency of the Duffy-null mutation due to selection 25-35 kya years ago in the western part of Africa, a geographical region and a time frame overlapping with the rise of another protective mutation, *β*^S^, the sickle-cell mutation protective at heterozygous state against the malaria caused by P. *falciparum*. In addition, the pattern of Duffy-null haplotypes highlights a quick spread of the Duffy-null allele in sub-Saharan Africa due to post-admixture selection events following the road of the recent Bantu expansion. Independent lines of evidence describing malaria as a life-threatening disease in West Africa from ∼30 kya, together with a rise in frequency followed by recent disseminations of the Duffy-null resistance, open new perspectives about both the history of malaria as a major human disease and the history of the main protective mutations in Africa.

## INTRODUCTION

Malaria is probably among the strongest, and the most documented, selective pressure imposed by an infectious agent on the human population (Kwiatkowski 2005). The burden imposed by malaria worldwide is predominant, with hundreds of thousands of deaths per year. In this context, balancing selection at the β-globin gene HBB, owing to the heterozygote advantage afforded by the hemoglobin β^S^ sickle mutation (Allison 1954; Allison 1956; Haldane et al. 2006), is the most iconic case of natural selection maintaining a protective mutation (at heterozygous state) against malaria caused by Plasmodium *falciparum*. The β^S^ mutation is widely distributed across sub-Saharan regions, with 9% and 7% average frequencies across agriculturalist and hunter-gatherer populations (Shriner and Rotimi 2018; Laval et al. 2019). Another mutation (T-42C, rs2814778), widely distributed across sub-Saharan regions at frequencies near fixation in Western, Central and Eastern African populations (Howes et al. 2011; McManus et al. 2017) was found to be protective against malaria caused by Plasmodium *vivax* (Wilairatana et al. 2022). The P. vivax malaria is the most geographically widespread human malaria with an estimated 4–5 million clinical cases per year, a strong heterogeneity in relapse risk between individuals (Stadler et al. 2022), and a particularly large hidden reservoir of asymptomatic.

The T-42C, rs2814778 mutation also known as the Duffy-null allele, FY*O or FY*B^ES^, is located in the transcription factor binding site in the DARC (Duffy antigen receptor for chemokines) promotor region. FY*B^ES^ may prevent vivax infections by causing erythroid-specific suppression of the expression of the DARC gene, also known as ACKR1 (atypical chemokine receptor 1) gene, a transmembrane receptor used by P. *vivax* to infect red blood cells (Höher et al. 2018). At the blood stage, vivax shows a strict preference for young CD71+ reticulocytes and classically uses the Plasmodium vivax Duffy-binding protein (PvDBP) to engage the DARC receptor on their surface, a pathway that has long been considered essential for invasion. At the molecular level, the FY*B^ES^ allele disrupts a GATA-1 transcription factor binding site in the DARC promoter, leading to an erythroid-specific silencing of DARC, while preserving its expression on endothelial cells. This selective loss of DARC abolishes the main receptor for the PvDBP on reticulocytes and underlies the near-complete protection of Duffy-negative individuals against classical P. vivax blood-stage invasion. Nevertheless, recent work indicates that rare infections can still occur in Duffy-negative hosts via low-level or transient DARC expression on erythroid precursors and potential alternative invasion pathways, suggesting that Duffy-null strongly reduces, but does not absolutely eliminate, the risk of vivax malaria (Shaikh et al. 2025). The role that natural selection for resistance to malaria may have played in the maintenance of high sub-Saharan frequencies of the Duffy-null allele is more controversial than for the β^S^ mutation. Despite being the most widespread of the malaria species, P. vivax is almost absent from Africa while the protective Duffy-null allele is almost absent in vivax endemic regions.

A study of the population of Grande Comore (Njazidja), living in a region with documented circulation of vivax and showing very high frequencies (0.86) of the Duffy-negative Fy(a–b–) phenotype due to a substantial sub-Saharan African ancestry, have suggested that African-derived FY*B^ES^ alleles could have experienced post-admixture selection in a vivax-exposed environment (Chiaroni et al. 2004). Several more recent studies conducted in Sahel, Madagascar and Pakistan where P. *vivax* also coexists with FY*B^ES^ brought by the sub-Saharan ancestry of the populations living in these regions, have clearly shown that selective pressures exerted by vivax have caused recent increase in Duffy-null frequencies (Hodgson et al. 2014; Triska et al. 2015; Laso-Jadart et al. 2017; Pierron et al. 2018; Cuadros-Espinoza et al. 2022). Based on intensive computer simulations and genomic data in Malagasi populations originating from admixture between Asian and African populations occurred ∼27 generations ago, two studies have shown that the high FY*B^ES^ frequency in Madagascar is explained by strong positive natural selection, with estimated selection coefficient (*s*), a proxy of the strength of selection, ranging from 0.066 to >0.2 respectively (Hodgson et al. 2014; Pierron et al. 2018). More recently, a study by Kasianov and colleagues showed post-admixture selection promoting the FY*B^ES^ introgression from Bantu speaking populations into the Khwe from the lower Okavango River Basin, an African region where vivax was only recently found (Kasianov et al. 2025). Finally, a study conducted in African regions where vivax is currently absent have shown a genome wide signal of positive selection that explains the Duffy-null frequencies near fixation in Sub-Saharan populations (McManus et al. 2017), with an Approximate Bayesian Computation (ABC) (Beaumont et al. 2002; Beaumont and Rannala 2004) estimation of the selection coefficient equal to 0.043 (95% CI: [0.011-0.18]). McManus and colleagues also estimated a start time of selection (*T*) 42,183 years ago (ya) (95% CI: [34,100-49,030]) for the youngest age found, based on the MRCA (more recent common ancestor) age of the FY*B^ES^ haplotypes. Assuming selection on standing variation with very low frequency of the Duffy-null allele at the onset of selection (𝑝^_!_ = 0.001), the study concluded to an ancient sweep, likely tens of thousands of years older than most other mutations associated with malaria resistance (McManus et al. 2017).

Yet, if this ancient sweep model may clearly explain the absence of vivax in sub-Saharan Africa by a progressive purge of the pathogen due to a quasi-fixation of Duffy-null and the resulting high rates ( >90%) of protected carriers, this model poorly explains the lack of this allele in Europe, or in Asia where the pathogen is still currently observed. The frequencies below 5% observed in Mediterranean populations are likely explained by recent migrations from Northern African and/or Middle East where Duffy-null is currently observed at frequency above 10%. Such a total, or quasi total, lack of FY*B^ES^ outside Africa is unlikely to occur assuming an ancient sweep with an onset of selection in a time frame overlapping with the estimated ages of the Out-of-Arica dispersal of modern humans into Eurasia, 40-60 kya (Gravel et al. 2011; Malaspinas et al. 2016; Mallick et al. 2016; Bergstrom et al. 2020; Choin et al. 2021; Ragsdale et al. 2023; Sümer et al. 2025).

Simulation-based ABC leveraging both computer simulations of the population history and various statistics computed from population genetics data, such as iHS developed to identify abnormally long haplotypes due to recent selection (Voight et al. 2006), have been proved to be efficient in estimating the age (*T*) of incomplete sweeps with current selected allele frequencies below 90%. For example, ABC provided consistent estimates of the age of selection targeting a non-synonymous variant (rs3827760, V370A) in the EDAR gene (𝑇 = 3,000[1,400 − 6,900]ya), or targeting the lactase persistence variant (rs4988235, MCM6-LCT region) (𝑇 = 7,441[6,256 − 8,683]ya) (Itan et al. 2009; Peter et al. 2012; Deschamps et al. 2016). As the inference of selection parameters can be influenced by past demography (Schraiber et al. 2016), simulation-based ABC are convenient for accounting for the complexity of past human demography inferred (Gravel et al. 2011; Schlebusch et al. 2012; Mallick et al. 2016; Schlebusch et al. 2017; Lopez et al. 2018; Bergstrom et al. 2020; Ragsdale et al. 2023). McManus and colleagues used such a demography-aware ABC (Peter et al. 2012) to jointly estimate the strength of selection (*s*) and the frequency *p*0 of the Duffy-null allele at the beginning of the sweep. To date, it is however unclear if existing ABC can provide consistent estimates of the age of complete sweeps, because selected allele frequencies at, or near to, fixation may erase the selection signal in surrounding haplotypes.

Here, we revisited the Duffy-null evolutionary history based on high coverage sequence data from 589 individuals distributed in 23 sub-Saharan populations of various geographical origins and modes of subsistence. We first evaluated from computer simulations the ancient sweep model assumed in Africa (McManus et al. 2017) and found that ancient selection ranging from 40kya to 70Kya poorly reproduces the Duffy-null frequencies observed world-wide, i.e. quasi fixed in Western sub-Saharan Africa and quasi absent in Eurasia. Seeking to estimate ages of selection that can better explain the lack of Duffy-null outside Africa, we first implemented a demography-aware ABC, calibrated to leverage the genetic information 200kb around the FY*B^ES^ mutation, and confirmed from simulation-based cross validations that such approaches poorly estimate the age and intensity of complete sweeps. To improve the joint estimations of *T*, *s* and *p*0, we implemented several supervised Deep Learning algorithms. We first implemented a convolutional neural network (CNN), a class of neural network successfully applied to infer population genetic parameters (Chan et al. 2018; Kern and Schrider 2018; Flagel et al. 2019) including selection (Torada et al. 2019), and potentially more efficient to leverage information contained in data (Laval et al. 2024). We found that the CNN, calibrated to leverage fine-grained features hidden in 1Mb of input sequence data around FY*B^ES^, may improve the estimations of *T*. To train this CNN with realistic simulated data reproducing the world-wide Duffy-null frequencies, we also implemented a generative adversarial network (GAN). Previous GANs using a generator producing genetic data from computer simulations and a neuron network as a discriminator that identifies simulated datasets indistinguishable from real ones, have been proved to be efficient in the prediction of demography and selection parameters (Wang et al. 2021; Riley et al. 2024). Under a composite past demography assuming a dispersal of modern humans from East Africa 50kya (Lopez et al. 2018; Ragsdale et al. 2023), our GAN and CNN predictions showed a rise in Duffy-null frequency due to selection 𝑇 = ∼30,000[22,000 − 37,000]ya, and located in the Western African lineage. This more recent time frame for selection than previously found can much better explain the quasi-total lack of Duffy-null outside Africa, and is, interestingly, compatible with the rise of β^S^, the protective mutation against the malaria caused by P. *falciparum*. Our study provides independent lines of evidence from another protective mutation and a different form of pathogen, that also identify malaria as a life-threatening human disease in Western Africa 20-30 kya. Finally, we found highly significant excesses in numbers of differences between Duffy-null and non-Duffy-null haplotypes, suggesting that high Duffy-null frequencies in Eastern African populations are due to post-admixture selection from a very recent admixture with newly arrived Bantu speaking populations.

## MATERIALS AND METHODS

### Ancient and modern data

We used genome-wide sequence data (30x) of 589 African individuals from various geographical location and public datasets, the Simons Genome Diversity Project (Mallick et al. 2016) and the GabonDiv datasets (Patin et al. 2017) (Table S1). We also used (the genotypes of the rs2814778 mutation of) 12 and 503 Europeans from the Simons Genome Diversity Project and the 1000 Genomes (1000G) project data (Auton et al. 2015), and 504 East Asians 1000G individuals. This dataset is divided in 34 populations: 6 populations of Western Bantu speakers (wBSP), 4 populations of Eastern Bantu speakers (eBSP), 4 populations of Western African agriculturalists (wAFR), 3 populations of Eastern African agro-pastoralists (eAFR), 2 populations of Western rainforest hunter-gatherers (wRHG), 2 populations of Eastern rainforest hunter-gatherers (eRHG), 2 populations of Southern hunter-gatherers (SHG), 6 populations of Europeans (EUR) and 5 populations of East Asians (ASI) (Tables S1). In addition, we used 19 ancient African individuals with radiocarbon ages ranged from 300 to 8,000 ya, and genotyped for rs2814778, ‘1240k capture’ Allele dataset (Mathieson et al. 2015) retrieved from the V44.3: January 2021 release at (Key *et al*. 2016; Lindo *et al*. 2016; Mathieson *et al*. 2018; Mathieson 2020; Ju and Mathieson 2021; Childebayeva *et al*. 2022; Kerner *et al*. 2023). All ancient individuals were treated as pseudo-haploids, i.e., hemizygotes for either the reference or the alternative allele.

### Neutrality statistics

We used the selink software (https://github.com/h-e-g/selink) to compute five neutrality statistics genome-wide. For each SNP we computed the matrix of pair-wise *F*ST (Weir and Cockerham 1984), a statistic reflecting the allele frequency differences between populations. For each SNP and in each population, we also computed 4 haplotype-based neutrality statistics, the iHS (Voight et al. 2006), the DIND (Barreiro et al. 2009), the ΔiHH (Grossman et al. 2010) and the nSL (Ferrer-Admetlla et al. 2014), which compare the haplotypes carrying the derived and ancestral alleles (Figure S1). To compute these haplotype-based neutrality statistics we used a sliding-windows approach with 100kb windows centered on each SNP (Fagny et al. 2014). The sliding-windows began and ended 50kb from the edges of each chromosome, to prevent early truncations in windows close to the edges. A similar approach was applied to the simulated regions used for simulation-based inferences described in the next sections. As iHS, DIND, ΔiHH and nSL are sensitive to the inferred ancestral/derived state, they were computed only when the derived states of SNPs were determined unambiguously, and were then normalized by DAF bin (Fagny et al. 2014; Voight et al. 2006) (mutations grouped by DAF bin, from 0 to 1, in increments of 0.025). In both empirical and simulated data, we applied classic filters by excluding minor alleles frequencies below 0.05, and we minimized the false-positive discovery by excluding SNPs with a DAF below 0.2, as the power to detect positive selection has been shown to be reduced at such low frequencies (Fagny et al. 2014; Voight et al. 2006). Finally, for each neutrality statistic, we defined the candidate SNPs of selection as being the top 1% (P<0.01) of SNPs genome-wide in real data, and simulation-wide in 100,000 neutral simulations.

### ABC estimations of T, s and ***p*0**

To jointly estimate, T, *s* and *p*0, we implemented a simulation-based ABC similar to the method previously applied to the Duffy-null allele (Peter et al. 2012; McManus et al. 2017). Because the haplotype-based statistics iHS, DIND, ΔiHH and nSL cannot be computed for fixed mutations such as FY*B^ES^ in many sub-Saharan populations, we incorporated linked variants around. In real data we selected a genomic region around the Duffy B variant (rs2814778). In computer simulations, we simulated genomic regions with a fake Duffy B variant located in the middle, and values of *T*, *s* and *p*0 randomly from flat prior distributions (Table S2). For every African population analyzed, each genomic region in real or simulated data was then summarized by the proportion of candidate SNPs determined separately for the *F*ST, iHS, DIND, ΔiHH and nSL. Note that the *F*ST mentioned is the pair-wise *F*ST between the analyzed population and a non-African sample, here the CEU population.

Concretely, empirical and simulated data were summarized by vectors of proportions of candidate SNPs computed for each neutrality statistic around the rs2814778. These vectors of summary statistics were ultimately used to fit simulated with empirical data, according to the standard ABC approach (‘abc’ R package, method = “Loclinear”) (Csillery et al. 2012). For ABC, we adopted the terminology classically used in machine learning. For convenience, the process of ABC model fitting, which involves determining the lowest ABC distances between simulated and empirical data, was abusively denoted as ‘training’, even though this step does not constitute a formal training as seen in typical machine learning approaches. ABC posterior distributions, point estimates (i.e., posterior average) and the 95% CIs were computed from the parameters underlying the 1,000 simulated data best fitting the empirical data.

### Convolutional Neural Network implementation

We implemented a Bayesian CNN algorithm with keras (Tensorflow as backdoor) to jointly estimate, T, *s* and *p*0 from sequence data around the Duffy B variant (rs2814778). The CNN also uses linked variants around FY*B^ES^, and the same neutrality statistics as in ABC, the *F*ST, iHS, DIND, ΔiHH and nSL (see above). Similarly to ABC the CNN also uses the proportion of candidate SNPs as statistics informative for selection. In contrast to ABC, the CNN does not summarize this information by computing each proportion once from all SNPs in the surrounding genomic region. Instead, the CNN is feed with 1D grey-scale images encoded from the proportions of candidate SNPs computed in 10kb sliding-windows centered on each SNP in the genomic region surrounding FY*B^ES^.

In contrast to fully connected neural networks, the CNN consists of consecutive sets of hidden neuron layers of different types, the convolutional and pooling layers, followed by a set of fully connected layers that carry out the image classification (Korfmann et al. 2023). CNNs are canonical feed forward neural networks with a first 1D convolution layer, which applies a convolution to the input image using a kernel (a filter or a matrix of learnable parameters (Korfmann et al. 2023)) to produce a feature map, which is then fed to a 1D maxpooling layer. This layer is used to reduce the dimensions of the feature map and capture coarse-grained features (features dissociated from their positions in the input image). We added three additional 1D convolutions and a last 1D maxpooling layer to capture more fine-grained features of the input image. A flatten layer is used to convert the feature map into a vector that is fed to several fully connected (dense) layers. The CNNs used in this study is inspired by the popular AlexNet model (Krizhevsky et al. 2017). During training, the CNNs parameters are adjusted with backpropagation of the gradient of the cross-entropy loss of function. Three output layers of 100 neurons each are used to obtain joint estimations of *s*, *T* and *p*0 through the softmax function, providing the likelihood of each parameter value/bin (parameters were discretized in 100 bins each; arithmetic precision at the second digit for floats). Posteriors averages were used as point estimates and the 95%CIs were computed from the posterior distributions obtained by resampling in the prior and likelihood distributions (Torada et al. 2019) The architecture, i.e., the numbers of layers, neurons, hyper-parameters (numbers of kernels and their dimensions) and the activation functions, are given in the Figure S2.

### Generative adversarial networks implementation

To train the CNN using data compatible with the world-wide pattern of Duffy-null frequencies, we implemented a generative adversarial network (using keras, Tensorflow as backdoor) able to closely reproduce real data (Figure S3). This GAN, inspired by efficient GANs previously developed (Wang et al. 2021; Riley et al. 2024), consists in a generator (population genetic data simulator) producing Duffy-null allele frequencies in and outside Africa, and a discriminator (multiplayer perceptron) that predicts whether the produced dataset is real or fake. The GAN implemented in this study uses a SLiM-based generator to generate frequencies according to the given demographic models described below. During training, the generator randomly draws parameters to use for the simulation software SLiM, and creates “fake” frequencies in Western African, European and East Asian populations (Tables S1). With feedback from the discriminator, the generator learns which parameters will produce the most realistic simulated data. The GAN-discriminator is a fully connected neural networks with 2 hidden layers. This multi-layer perceptron (MLP) is a feed forward network feed with the vector of populations frequencies, and corresponding pair-wise *F*ST if necessary. A single output layer of 2 neurons is used to obtain the state of the input data through the softmax function, providing the likelihood of each state, “fake” (0) or “true” (1). As in Riley and colleagues, we trained the generator using a gradient-free approach. The GAN-discriminator is trained once for all using real data labelled “true” (1), complemented with simulated data labelled “fake” (0), which exhibit a world-wide frequency pattern different to that observed for Duffy-null. During training, the parameters of the MLP are adjusted with backpropagation of the gradient of the cross-entropy loss of function. The GAN-generator is trained using computer simulations and a rejection/acceptation algorithm, which accepts the parameter values that produces data declared to be “true” by the GAN-discriminator. At the end of training the GAN-generator learnt the probability distribution of each parameter maximizing the probability to generate a realistic world-wide pattern of Duffy-null frequencies.

### Training the GAN, ABC and CNN algorithms

For each analyzed population, the ABC, GAN and CNN are trained using 1Mb of simulated sequenced data, with a fake Duffy B variant (rs2814778) located in the middle. The GAN is trained with the simulated Duffy-null frequencies averaged across 10 Western African, 6 European and 5 East Asian populations mimicking our empirical samples (Tables S1), and with the corresponding matrix of average *F*ST. ABC and CNN are trained with the proportions of candidate SNPs determined for 5 neutrality statistics separately, *F*ST, iHS, DIND, ΔiHH and nSL (see above). ABC is trained with the proportions of candidate SNPs computed in 100kb and 200kb around the simulated Duffy B variant.

The CNN is trained with the proportions of candidate SNPs computed 10kb around 500 randomly drawn SNPs linked to the simulated Duffy B variant. Concretely, GAN and ABC use vectors of statistics of dimensions (1 × 6) and (1 × 10=5×2) respectively, and CNN uses vectors of dimension (1 × 2500=5×500) encoded into 1D dimension images in which the 2500 pixels exhibit values ranged from 0 (no candidate for selection 100kb around) or 1 (all SNPs are candidate for selection 100kb around). Each vector and 1D image was simulated to match the number of populations, sample sizes and geographical locations of our samples, under a specified demographic model and values of *T*, *s* and *p*0, drawn from known prior distributions presented in the next section below. Finally, the resulting simulated vectors labelled with the corresponding *T*, *s* and *p*0 values are ready to use for training in a supervised learning. The GAN and ABC are trained using 1,000,000 and 100,000 simulated vectors. The CNN is trained using 100,000 simulated 1D images during a rather short number of 3 epochs. The accuracy of the GAN, ABC and CNN predictions were evaluated using a classical cross validation procedure performed based on 200 simulated data not used during the training process (these simulated data, used as empirical data for which the true values of *T*, *s* and *p*0 are known, are referred to as pseudo-empirical datasets in the following sections). Note that the corresponding empirical vectors and images used for GAN, ABC and CNN predictions were obtained of the same manner and are of the same dimensions as those used during trainings.

### Forward-in-time computer simulations used to produce the training datasets

Sequence data were simulated with SLiM (Haller and Messer 2017; Haller and Messer 2019), using randomly drawn human compatible recombination maps (Auton et al. 2015), and under various past demography incorporating divergence times, migration rates, population admixture and reduction/growth of effective sizes for 34 populations: 6 wBSP, 4 eBSP, 4 wAFR, 3 wAFR, 2 wRHG, 2 eRHG, 2 SHG, 6 EUR and 5 ASI (Tables S1). Because such demographic parameters have never been jointly estimated for these 34 populations together, the simulated demographic model is a composite demography drawn from several studies (Figure 1, Table S2): split times between Bantu speakers and rainforest hunter gatherers were drawn from (Lopez et al. 2018), migration rates and split times between Southern hunter gatherers, Western and Eastern Africans and Eurasians populations were drawn from (Ragsdale et al. 2023). Connections between Western Bantu speakers and Western Africans were drawn from (Schlebusch et al. 2012; Schlebusch et al. 2017). In this composite model, the Out of Africa dispersal of modern humans originates from the Eastern African lineage (eAFR) 50kya. In parallel, we also simulated another version of our composite model but close to model used by McManus and colleagues (Gravel et al. 2011), which set an Out of Africa dispersal 50kya originating from the common ancestor of Western, including Bantus speakers, and Eastern Africans.

**Figure 1.**
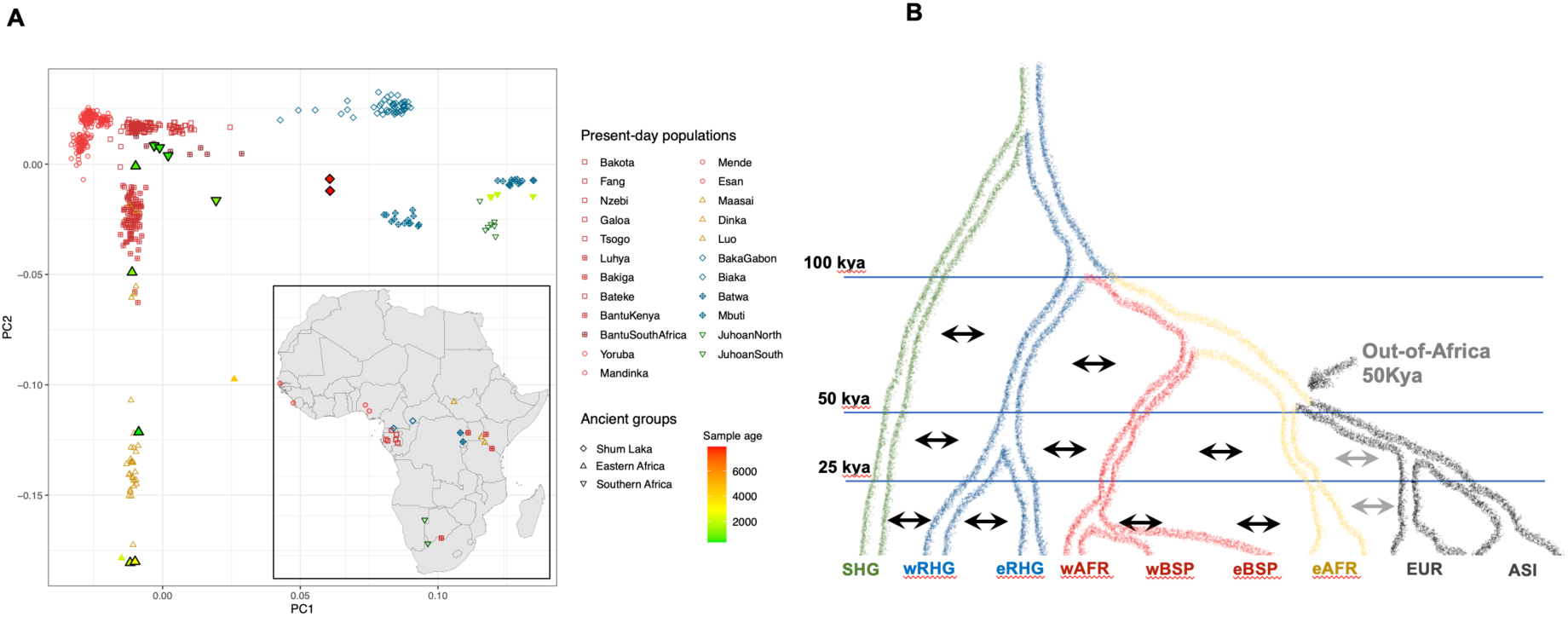
Sub-Saharan samples analyzed (A) PCA and geographical locations of the sus-Saharan populations included in the dataset. The corresponding sample sizes, the Duffy-null frequency, and the neutrality statistics computed for the Duffy Variant (*F*ST iHS, DIND, ΔiHH and nSL) are indicated in Table S1. (B) The composite past demography merging the Lopez’s model, for the BSP and RHG samples, and the Ragsdale’s model, for the East Africa dispersal, used to simulate 1Mb of sequence data in the 34 populations in the dataset: 6 wBSP, 4 eBSP, 4 wAFR, 3 wAFR, 2 wRHG, 2 eRHG, 2 SHG, 6 EUR and 5 ASI.

In this case the East West split in Africa was set to 45kya (Schlebusch et al. 2012). Because SLiM is a forward-in-time simulator, the computation times depend on both the effective population size *N* and the number of generations *t* considered. Effective population sizes and times were thus rescaled according to 𝑁_"_/𝜆 and 𝑡/𝜆, with 𝜆 = 10 (Hoggart et al. 2007; Haller and Messer 2017; Haller and Messer 2019). For each simulation, the age of selection *T*, the strength of selection as measured by the selection coefficient *s*, and the frequency of the Duffy-null allele at the onset of selection *p*0 were randomly sampled from uniform prior distributions: *T* ∼ *U*(0, 100,000) ya, *s* ∼ *U*(0.01, 0.1) under an additive model (*h* = 0.5), and *p0* ∼ *U*(1/*N*, 0.2).

### Simulation-based evaluation of the GAN, ABC and CNN algorithms

The performances of our GAN ABC and CNN algorithms were evaluated based on pseudo-empirical data, i.e., simulated data not used in the training steps but used as empirical data for which the true values parameters are known, as described above. These pseudo-empirical data were simulated using composite demographic models to reproduce the effects of drift, migration waves, recent admixture, and the features of the empirical data used in this study (number of populations and samples per population), as described in the section above. Based on these pseudo-empirical data, we then assessed the accuracy of methods tested by comparing predicted and simulated parameter values, 𝜃^<^_#_ and 𝜃_#_, respectively obtained for a pseudo-empirical data *p*. We used classic accuracy indices computed in cross-validation procedures: the linear correlation coefficient *r* computed between 𝜃^<^_#_ and 𝜃_#_, the relative root of the mean square error, *RMSE* (i.e. the root of the *MSE* expressed as a proportion of the true value, 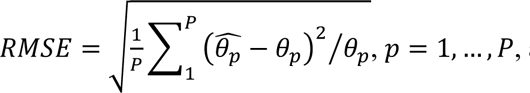, and the proportion of true values within the 90% credible intervals of estimates, 𝐶𝐼𝑐𝑜𝑣 = ^$^ ∑^%^ 1(𝑞_$_ < 𝜃_’_ < 𝑞_&_ ) where 1(*C*) is the indicative function (equal to 1 when *C* is true, 0 otherwise) and 𝑞_$_ and 𝑞_&,_the corresponding percentiles of the posterior distributions.

## RESULTS

### The current ancient sweep model poorly explains world-wide Duffy-null frequencies

To revisit the Duffy-null evolutionary history, we used high coverage sequence data in 589 individuals from various sub-Saharan locations, with 515 and 504 Europeans and Asians respectively (Auton et al. 2015; Mallick et al. 2016; Patin et al. 2017) (Figure 1A) (Table S1). These individuals are distributed in 34 populations, 23 in sub-Saharan Africa, 6 in Europe and 5 in East Asia. In the western part of Africa, the FY*B^ES^ variant was found fixed in every four populations of Western African agriculturalists (wAFR) and in every 10 populations of Bantu speakers (w/eBSP), except in the Bakiga (𝑝^ = 0.98) currently located in East Africa. Note that in this manuscript, the Eastern and Southern Bantu speakers (eBSP, sBSP) are considered to be members of the Western African lineage due to the very recent Bantu dispersal from west Africa in the last three millennia (Patin et al. 2017). The Duffy-null allele was also found fixed in every four populations of rainforest hunter-gatherers (w/eRHG), except in the Batwa (𝑝^ = 0.714) one of the two Eastern rainforest hunter-gatherers (eRHG) populations in the dataset. Note that in the two populations of Southern hunter gathers (SHG), where the Duffy-null allele is expected at intermediate values (∼0.3), the frequency observed equal to 0 can be explained by very low sample sizes (5 and 2 individuals respectively). To summarize, the Duffy-null allele was found fixed in all sub-Saharan Africans except in three populations living in the eastern part, including the Maasai (𝑝^ = 0.885), one of the three Eastern African agro-pastoralists (eAFR). As expected, the Duffy-null allele was found totally absent in East Asia and Europe, except in the two Mediterranean European 1000G populations included in the dataset (𝑝^ < 0.02), likely due to very recent migrations as already mentioned.

With a strong intensity of selection often exceeding s=0.066 and clearly demonstrated in admixed population living in the dispersal area of *vivax* (Hodgson et al. 2014; Triska et al. 2015; Laso-Jadart et al. 2017; Pierron et al. 2018; Cuadros-Espinoza et al. 2022), the common view about the evolutionary history of FY*B^ES^ assumes an ancient selective sweep from a very low frequency (𝑝R_!_ = 0.001) (McManus et al. 2017). We first aimed to evaluate the ancient sweep inferred; a strong selection (𝑠^ = 0.043[0.011 − 0.18]) starting around 𝑇 = 42,183[34,100 − 49,030]ya based on the major FY*B^ES^ haplotype (𝑇 = 56,052[38,927 − 75,073]ya based on the minor haplotype). We simulated this ancient sweep model under a composite past demography build for our 34 populations (Lopez et al. 2018; Ragsdale et al. 2023) and updated to incorporate a modern human dispersal 50kya from the common ancestor of Western and Eastern African lineages (Gravel et al. 2011) (Figure S4, Material and Methods). We found that an ancient sweep poorly reproduces the lack of Duffy-null in Europe and East Asia (Figure S5). For example, with an onset of selection simulated at 𝑇 = 42kya the Duffy-null variant was observed at high frequency in Europe and/or East Asia in 99.99% of computer simulations (Figure S5). These results, confirming that the total, or quasi total, absence of FY*B^ES^ outside Africa is unlikely to occur assuming a sweep starting in a time frame close to the Out-of-Arica dispersal, suggest that the Duffy-null selection parameters need to be confirmed by independent estimations, mainly for the age of selection.

Aiming to jointly estimate *T*, *s* and *p*0, we implemented a simulation-based ABC method based on the *F*ST and various haplotype-based statistics (Peter et al. 2012; McManus et al. 2017). Because iHS and related haplotype-based statistics cannot be computed for fixed mutations such as FY*B^ES^, our ABC also incorporates linked variants around. Specifically, the ABC uses the proportion of candidate SNPs of selection, computed for each statistic used (Material and Methods). The proportions of candidate SNPs of selection around a selected variant are helpful statistics to quantify selection by leveraging the hitchhiking effect of positive selection. Selective sweeps indeed produce clusters of candidate SNPs in the vicinity of selection targets, whereas under neutrality candidate SNPs are more uniformly scattered (Voight et al. 2006). Cross-validations based on pseudo-empirical data, i.e., simulated data used as empirical data for which the true values of selection parameters are known (Material and Methods), showed that ABC poorly estimates the age and the intensity of selection in case of complete or quasi complete sweeps (Figure S6). This result highlights the need for new methodologies to better leverage the genetic information hidden in the data. Finally, many studies have shown that unlike in the demography assumed for the ancient sweep model, Western and Eastern African lineages diverged prior to the out of Africa dispersal, which likely originates from the Eastern African lineage (Malaspinas et al. 2016; Schlebusch et al. 2017; Ragsdale et al. 2023). This also highlights the need to incorporate a better modelling of the past demography assuming a human dispersal from the Eastern part of Africa.

### The rise of the Duffy-null variant in the Western African lineage

To investigate the selection parameters *T*, *s* and *p0* underlying the Duffy-null evolutionary history we leveraged 1Mb of sequenced data around the Duffy-null variant obtained in the 23 in sub-Saharan African populations presented above. To perform the estimations, we used two deep learning algorithms. For improving the estimation of the age of selection, we applied a Bayesian convolutional neural network, which jointly infer the posterior distributions of *T*, *s* and *p*0 (Material and Methods). This CNN also uses candidate SNPs of selection (also determined for *F*ST, iHS, DIND, ΔiHH and nSL separately), but in contrast to ABC, the CNN leverages 1Mb of fine-grained genomic variation of the proportion of candidate SNPs around FY*B^ES^, encoded in 1D grey-scale images. In these 1D grey-scale images the pixel colors are the proportions of candidate SNPs in 10kb sliding-windows centered on each SNP around FY*B^ES^. Cross-validations based on pseudo-empirical data confirmed that CNN may improve the estimates of *T* (Figure S6). In addition, to estimate the selection parameters based on past human demography assuming a more realistic dispersal from the Eastern African lineage, we aimed to train our CNN with computer simulations under the dispersal parameters inferred by Ragdsdale and colleagues (Ragsdale et al. 2023) (Figure 1B, Material and Methods). Because we have shown that most computer simulations did not reproduce the world-wide Duffy-null frequencies, for instance more that 99% of simulations result in a quasi-fixation outside Africa under the ancient sweep model (Figure S5), we also aimed to avoid training our CNN with such high rates of unrealistic simulated data. We thus used another Deep Learning algorithm, a generative adversarial network (Wang et al. 2021; Riley et al. 2024) able to better reproduce the quasi total lack of Duffy-null outside Africa. The implemented GAN uses SLiM as a generator to produce fake data from computer simulations and a multi-layer perceptron as a discriminator that predicts whether the produced dataset is real or fake (Material and Methods). We cross-validated the GAN predictions and found that the trained GAN-discriminator is highly efficient in identifying simulated data reproducing both the fixation in West Africa and the lack of the Duffy-null allele in Europe and East Asia (less than 1% of error which are systematically made for simulated frequency ranged between 0.95 and 0.99 in western Africa, Figure S7).

Finally, before applying our CNN to estimate the selection parameters, we first applied our trained GAN-discriminator to determine if the rise in frequency of the Duffy-null mutation occurred either concomitantly in all geographical regions of Africa, or in a given specific sub-Saharan lineage. We thus simulated various scenarios under our composite past demography with a dispersal of modern humans from East Africa 50kya (Lopez et al. 2018; Ragsdale et al. 2023) (Figure 1B, Material and Methods), and randomly drawing parameters in specified flat priors, including *T*, *s* and *p0* (Table S2). We simulated several scenarios with an onset of selection, ranged from present day to 100,000 years in the past, located either in the RHG lineage (wRHG, eRHG), in the Eastern African lineage (eAFR) or in the Western African lineage (wAFR, wBSP, eBSP and sBSP). We found that the percentage of simulations declared as “true data” by our GAN-discriminator in the Western African scenario is at least 40 times higher than in the other alternatives (Figure 2A). This large excess of simulated data reproducing both the fixation in West Africa and the lack of the Duffy-null in Eurasia, result in Bayes Factors higher than 40, strongly supporting the western scenario (𝐵𝐹 > 40). According to the Jeffreys’ scale, the GAN-discriminator provides very strong strength of evidence for the rise of the Duffy-null allele due to positive selection in the western part of Africa.

**Figure 2.**
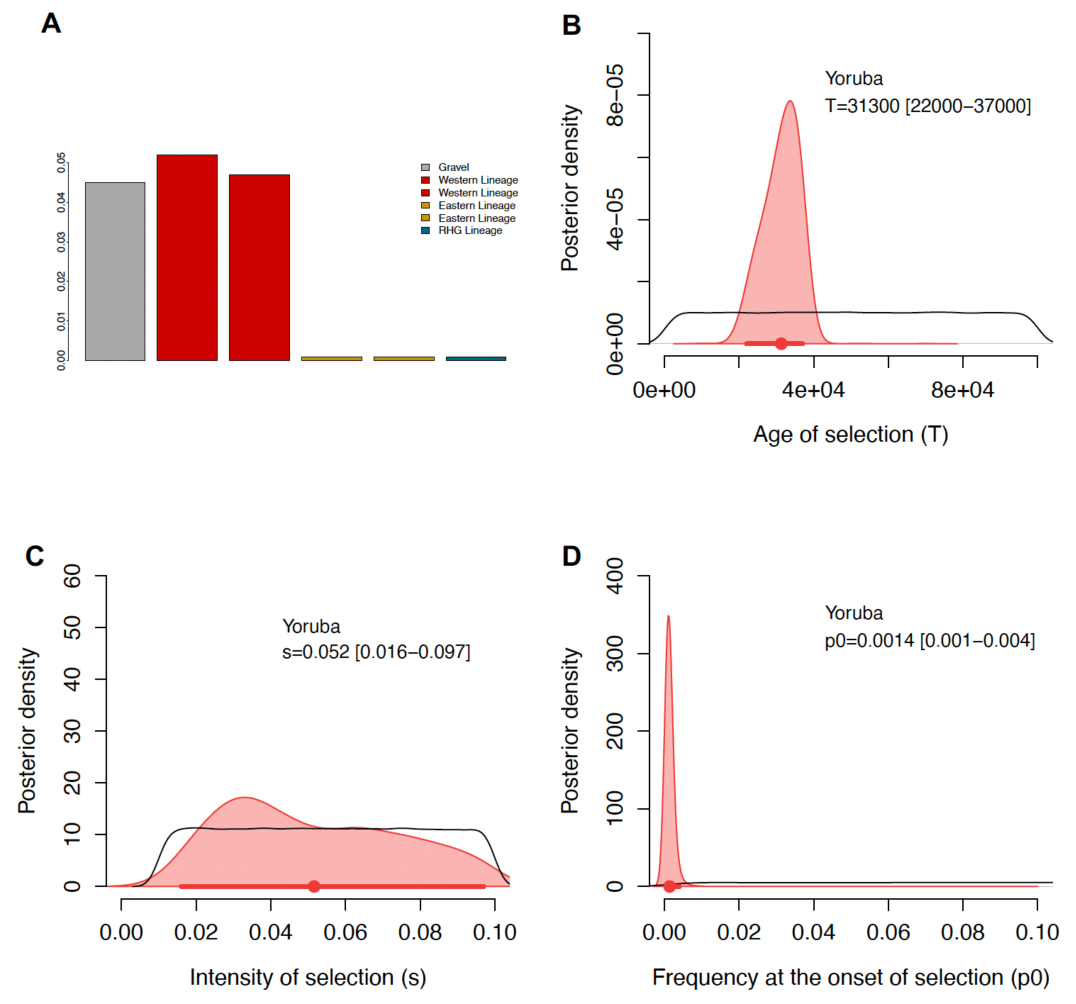
GAN predictions and CNN estimations of the selection parameters *T*, *s* and *p*0 (A) Percentages of computer simulations declared as “true data” by our GAN-discriminator (Figure S3), i. e., percentages of simulations reproducing both the fixation of Duffy-null allele in West Africa and the lack of this allele in Europe and East Asia. The simulations were performed under the composite model displayed in Figure 1B, and using the prior distributions indicated in Table S2, *T* ∼ *U*(0, 100,000) ya, *s* ∼ *U*(0.01, 0.1) under an additive model (*h* = 0.5), and *p*0 ∼ *U*(1/*N*, 0.2). 100,000 simulations were performed in each simulated scenario: onset of selection in the Western African lineage (red), in the Eastern African lineage (yellow) the RHG lineage (bleu). Note that we show two sets of 100,000 simulations for the western and eastern scenarios. We also included simulations performed under the composite model dispersal from the ancestor of Western and Eastern Africans (grey) (Figure S4). (B-D) Posterior distributions obtained with our CNN algorithm in the Yoruba population (sample size of 76 individuals) for *T* , *s* and *p*0 respectively. The black lines show the prior distributions. Point estimates (posterior mean) and 95% confidence interval (in brackets) are indicated and displayed with a dot and bold line in the bottom of the posteriors.

### CNN predicts an onset of selection 30k years ago

Based on a scenario of the rise of FY*B^ES^ clearly established in the western part of Africa, we jointly estimated *T*, *s* and *p*0, performing as follow. We first trained the GAN using successive learning cycles (Figure S3). The GAN-generator simulates the onset of selection in the Western African linage as previously performed; under our composite past demography with a dispersal from East Africa and drawing parameters in specified flat priors (Table S2). The simulation is presented to the GAN-discriminator and the underlying parameters are accepted if the discriminator predicts a “true data”, rejected otherwise. This learning circle is repeated 1,000,000 times to properly explore the parameter space. Once these one million leaning circles are over the training is completed, the GAN is ready, the GAN-generator has learnt the distribution of parameters generating realistic data with both fixation and absence of FY*B^ES^ in and out Africa respectively. We then used the trained GAN-generator to simulate 100,000 computer simulations of 1Mbp of sequence with a fake selected mutation in the middle. These simulations were used to train our CNN algorithm and obtain the CNN predictions in real populations. The posterior distributions of *T*, *s* and *p0* obtained by the trained CNN applied to the Yoruba population, the population with the highest sample size (72 individuals, Table S1), are shown in Figure 2B-D. While we found very similar intensity 𝑠^ = 0.052[0.016 − 0.097] and frequency at the onset of selection 𝑝R_!_ = 0.0014[0.001 − 0.004] as those found by McManus and colleagues (McManus et al. 2017), we estimated a much more recent onset of selection 𝑇 = 31,300[22,000 − 37,000], likely ten thousands of years younger than the 42kya previously found. Interestingly, the estimated onset of section in the Western African lineage was very similar independently of the populations used (Figure 3A, the corresponding posterior distributions of *s* and *p*0 are displayed in Figures S8,9). All these CNN predictions clearly show a rise in frequency of the Duffy-null allele due to an onset of selection 25-35kya in the western part of Africa. In addition, the very low frequency at the onset of selection (∼0.001) found both by McManus and colleagues and in this study indicates that, at this time, Duffy-null was a young but preexisting mutation occurred shortly before selection starts. This predicted onset of selection of ∼30kya on a preexisting protective variant therefore suggest that vivax malaria became a life-threatening human disease in Western Africa likely during this period.

**Figure 3.**
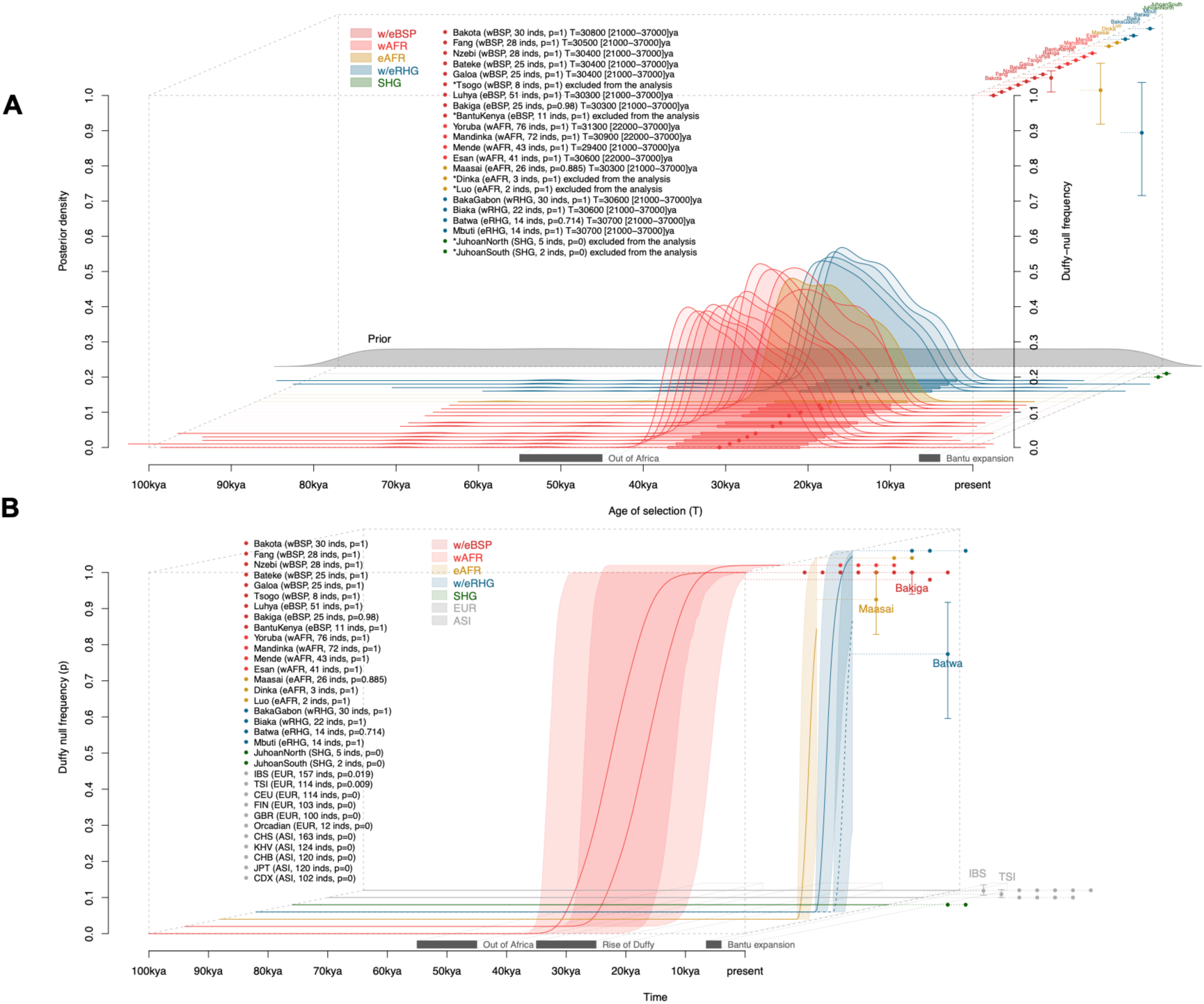
Duffy-null frequency trajectories during the last 100,000 years (A) Posterior distributions of the age of selection *T* obtained in every population analyzed. Posteriors are displayed in the order of appearance in the legend (first populations on the top). We excluded populations with a sample size lower than 14 individuals from the analysis (indicated by * in the legend). The grey surfaces show the prior distributions. Point estimates (posterior mean) and 95% confidence interval (in brackets) are also indicated in the legend and displayed with a dot and bold line in the bottom of the posteriors. The corresponding frequency of the Duffy-null allele are indicated in the legend and displayed on the right axis. (B) Simulated frequency trajectories during the last 100,000 years under our selection model inferred. Note that the bleu dashed line (and the corresponding blue surfaces show the trajectory in the eRHG lineage. The corresponding frequency of the Duffy-null allele are indicated in the legend and displayed on the right axis (displayed in order of appearance in the legend).

### Post-admixture selection of Duffy-null haplotypes in East Africa

Based on the GAN and CNN predictions, we set an updated selection model for the Duffy-null allele across the last 100,000 years, highlighting a sweep starting ∼30kya in the Western African lineage, with an intensity of selection ranged between 0.01 and 0.1 and a frequency at the onset of selection close to 0.001. To check the validity of this model we performed a posterior predicting validation analysis by checking if this model provides simulated data compatible with the observed Duffy-null frequencies. Unlike in the ancient sweep model where the proportion of compatible simulated data was found lower than 10^-2^ (Figure S5), we found that this selection model produces ∼25% of simulations declared to be compatible with real data by the trained GAN discriminator (these compatible simulations are shown in Figure 3B). This result shows that the parameters ranges predicted by CNN appear to describe a more realistic evolutionary history of the Duffy-null allele, which can explain both the fixation and the lack of Duffy-null in West Africa and Eurasia (Figure 3B, red and gray curves showing the simulated frequency trajectories across the last 100,000 years in Western African and Eurasian lineages respectively, the Southern hunter gatherers being excluded from the analysis due to very low sample sizes). It is worth mentioning that most frequency trajectories in West Africa exhibit high frequency from 20-10kya, which is compatible with the ancient data available in the region (two ancient individuals sampled in Shum Laka, Cameroon, are Duffy-null carriers ∼8,000 ya, Table S1).

However, this model did not reproduce the fixed frequencies in the RHG and Eastern African lineages respectively. Assuming a commonly inferred gene flow of the order of 10^-4^ and 10^-5^ within and between continents (Ragsdale et al. 2023) (Table S2), the Duffy-null allele necessarily spreads by migration out of the Western African lineage, invading all the other populations, including Europeans and East Asians, where it quickly increases in frequency due to selection. Consequently, in most computer simulations reproducing the absence of Duffy-null out of Africa, the allele did not migrate out of the Western African lineage. We assumed that high Duffy-null frequencies currently observed in eAFR and eRHG (Figure 3B) are due to recent admixture with the Bantu speaking populations arrived in Eastern Africa the last millennia. To confirm this assumption, we compared the Duffy-null haplotypes observed in the Maasai and Batwa populations with those observed in Luhya, the eBSP population with the largest sample size, used here as a putative source population. We used the average numbers of differences between pairs of haplotypes: between carrier and non-carrier of Duffy-null, Π_)*+,-,)*_, and between Duffy-null haplotypes, Π_)*+)*_ (computations made excluding Duffy-null and old variants with frequency higher than 0.5). We found highly significant excesses in numbers of differences between Duffy-null and non-Duffy-null haplotypes, both within and between populations. For instance, using 10kb haplotypes centered on the Duffy-null mutation, Π_)*+,-,)*_within the Maasai and Batwa populations are at least twice higher than the corresponding Π_)*+)*_ (𝑃 = 1.7 10^+./^ and 𝑃 = 1.18 10^+0$^respectively, Table S3). Such significant excesses of pairwise differences suggest that Duffy-null is unlikely to be a mutation occurred in an East African haplotypic context. In addition, the Duffy-null haplotypes observed in Massai, Batwa or even in Mbuti are indistinguishable from those observed in Luhya (Π_)*+)*_between these populations and Luhya are very similar to Π_)*+)*_within Luhya, ∼2 differences between pairs of haplotypes in average Table S3, Figures S10-14). These results indicate that the high frequencies of Duffy-null haplotypes in East Africa, including the Eastern rainforest hunter-gatherers, are likely due to post-admixture selection from a recent admixture with one or several newly arrived Bantu source populations. Beyond the younger age of selection inferred, our study also highlights an evolutionary history with a quick spread of the Duffy-null allele in sub-Saharan Africa due to post-admixture selection events along the road of the recent Bantu expansion, as also indicated by recent results obtained in the Khwe from the lower Okavango River Basin (Kasianov et al. 2025).

## DISCUSSION

Here, we present demography-aware Deep Learning algorithms designed for the estimation of key selection parameters underlying selective sweeps. The GAN was trained to be applied on variants evolving under population specific complete sweeps, i.e. both fixed in a population and absent elsewhere, a scarce situation in humans but likely more prevalent in other species such as Drosophila (Garud et al. 2015; Garud et al. 2021). The CNN applied to complete sweeps showed a moderate gain in accuracy compared to an equivalent ABC, as shown by our cross validations performed assuming *p0* of the order of 0.001, which is “in fine” the estimated values of the Duffy-null allele. The CNN can however be applied on incomplete sweeps with potential gains in accuracy which should be assessed regarding the results obtained in previous implementations of CNNs in the estimation of selection parameters (Torada et al. 2019).

Our study estimates more recent ages of selection, 𝑇^)^∼30kya, ∼10 thousand years younger than previously thought, that are much compatible with the lack of Duffy-null in endemic regions of vivax. This age should not be interpreted as the age of the mutation (date of the occurrence in populations), since Duffy-null could be present in populations before the beginning of selection. However, our estimations of a very low frequency when selection started, 𝑝^_!_∼0.001, also informs that FY*B^ES^ is likely a mutation occurred shortly before selection. Predicted *p*0 are proxy of the age of mutations at the onset of selection because frequencies of neutral mutations correlate with their age. Basically, under the Duffy-null selection model, older mutations with their corresponding higher *p*0 are excluded because they may predate the split with the eastern lineage, then migrate out of Africa, and thus be also present at high frequency in endemic regions of vivax (older mutations with *p*0>0.01 are expected at high frequency in Europe and/or East Asia in our simulations). The estimations of low *p*0 also suggest that FY*B^ES^ likely occurred in the Western African lineage, since a neutral mutation arrived by gene flow is expected at intermediate frequency in the source population. Such a mutation may likely predate the out of Africa dispersal and be also present at high frequency in Eurasia.

Our predictions were made under the admitted assumption of positive selection driving the evolutionary history of Duffy-null, as clearly indicated by many studies conducted in admixed populations coexisting with vivax. Our predictions also confirm that vivax has been present in Africa for several tens of thousands of years, since ∼30kya at least. The alternative assumption of vivax totally absent from Africa until a very recent arrival in region (Kasianov et al. 2025) is not supported by the data. Again, in absence of the vivax selective pressures, a neutral Duffy-null mutation nearly at fixation must be an old mutation, predating both the split between African lineages and the out of Africa dispersal, which must be also observed out of Africa. The only valid assumption without vivax in Africa is the past existence and disappearance of an unknown African plasmodium also causing life-threatening malaria and Duffy-dependent to infect human erythrocytes. Only further biological studies showing evidence for Duffy-dependent pathways, for instance in ancestors or relatives of falciparum, can help to test such alternative hypotheses.

Our study also tells us that 30kya vivax malaria became a significant selective pressure in West Africa. Interestingly, the rise in frequency of the β^S^ sickle mutation estimated in an overlapping time frame (∼22 [10-50]kya) (Laval et al. 2019) also suggested that falciparum malaria became a significant selective pressure in the same period. Two independent mutations point toward a similar period for the rise of malaria caused by two independent pathogens, suggesting that around 20-30kya the epidemiologic parameters or the environment of Western African populations likely changed and promoted the spread of plasmodium (increase of population density, deforestation). The genetic diversity of present days African populations are compatible with population growth 16,000–22,000 ya (Patin et al. 2009; Patin et al. 2014), suggesting that increased population density could facilitate the transmission of plasmodium in populations. We can notice that only ∼10% of the sub-Saharan Africans are protected carriers against falciparum by the β^S^ mutation (heterozygotes β^S^ carriers) while nearly all are protected carriers against vivax due to the Duffy-null mutation. Competition between vivax and falciparum together with a large discrepancy between rates of protected carriers in the host populations, a factor that can potentially impact the fitness of pathogens, may partly explain why vivax, with a lower fitness, was eliminated while falciparum, with a higher fitness, is still so prevalent.

Under the assumption of commonly admitted migration rates within and between continents, only a “migration + recent Bantu admixture” gene flow model can reproduce the pattern of Duffy-null frequency in East Africa, highlighting the role of recent admixture in the spread of advantageous mutations in humans. The frequency trajectories under this modeling in East Africa reproduced high FY*B^ES^ frequencies thanks to a post-admixture selection from recently arrived Bantu populations (see the eRHG and eAFR, displayed in yellow and dashed blue lines in Figure 3B). Trajectories with high frequency ∼2,000 ya in Eastern African lineages are compatible with ancient data (ancient individuals in Kenya are Duffy-null carriers ∼2,800 ya, Table S1). The analyses based on iHS were inconclusive in Massai and Batwa despite present day frequencies of 0.885 and 0.714 respectively (Table S1), likely because of a lack of iHS power excepted under selection on high standing variation due to recent admixture. In contrast, a fine analysis of Duffy-null haplotypes confirmed the assumption that high frequencies observed in our sub-Saharan African populations may result from post-admixture selection from Bantu speaking populations. Note that, no “migration + recent admixture” model can reproduce the data if selection starts in the Eastern African lineage. First, there are no known recent east-west migration waves of comparable magnitude with that of the Bantu expansion that could explain a recent arrival of Duffy-null in Western Africa. Second, there is no selection model of equal magnitude across Africa that could reproduce a selected mutation both totally fixed in the region very recently colonized while being at 0.7 frequency in the region where the selection started several thousand years before. The latter also excludes an origin of selection in Southern Africa where the Duffy-null frequency is currently below 0.5. This confirms again that Duffy-null likely started to rise in frequency due to positive selection in the western part of Africa.

However, an increasing number of reports of vivax malaria in genotypically Duffy-negative individuals, together with experimental work on alternative ligands such as PvRBP and PvEBP/DBP2, indicate that P. vivax can exploit additional Duffy-dependent or Duffy-independent routes to enter reticulocytes. These biological features – a large latent hypnozoite reservoir, highly reticulocyte-restricted invasion, and emerging plasticity in invasion pathways – help to explain both the resilience of P. vivax transmission and the strong, but complex, selection pressures it can impose on host loci such as DARC (Noviyanti et al. 2022). Beyond its role as a malaria receptor, DARC also functions as an atypical chemokine receptor that scavenges and presents multiple CC and CXC inflammatory chemokines, thereby modulating leukocyte trafficking and systemic inflammatory tone. Individuals carrying the Duffy-null allele frequently display so-called benign ethnic neutropenia, characterized by chronically low absolute neutrophil counts without increased risk of bacterial infection, an effect now clearly attributed to rs2814778 at DARC rather than to global African ancestry. Experimental and clinical data suggest that loss of erythroid DARC alters the chemokine “sink” and can reshape neutrophil production, mobilization and tissue homing, with context-dependent consequences on immune-mediated pathology and susceptibility to other infections such as HIV. These pleiotropic effects indicate that selection on FY*BES in Africa likely acted not only through protection against P. vivax, but also through a broader reshaping of host–pathogen and host–inflammation interactions (Bouyssou et al. 2022).

An apparently conflicting observation comes from the Jarawas (Das et al. 2005) a small relict population from the Andaman Islands, where serological typing reported 100% Duffy-negative phenotypes. At face value, such data could be interpreted as evidence that Duffy negativity was already present in early Out-of-Africa migrants. However, these observations rely exclusively on classical blood group serology and do not provide any information on the underlying molecular mechanism, nor on the presence of the canonical African FY*B^ES^ (T-42C, rs2814778) allele. In principle, complete Duffy negativity in Jarawas could reflect distinct regulatory or coding variants at DARC, undetected by serology, or even genetic drift acting on rare Duffy-negative alleles in a very small and isolated population. Without sequence-level data at the DARC locus, the Andaman findings therefore do not contradict a model in which FY*B^ES^ arose and was driven to high frequency specifically within Western African lineages after the main Out-of-Africa dispersal, but instead highlight the possibility of convergent or parallel routes to Duffy negativity in different human groups.

In conclusion, the GAN and CNN algorithms presented here offer alternative ways for analyzing populations specific complete sweeps. However, our approach does have some limitations. Our simulated models are dependent on our current limited knowledge of past migrations in humans.

However, our conclusions do not change when increasing the migration intensity; selection still starts in West Africa (the number of simulations without Duffy-null outside Africa diminish). Only models with total absence of migration, for instance no migration between East Africa and Eurasia during long periods, may alter our conclusions. Our simulated training datasets also assumes a constant selection coefficient (*s*) across time and geographical regions. These reasonable assumptions may be updated during GAN - CNNs training, but empirical results in Malagasy and more recently in Khwe (*s* ranged from 0.035 to 0.114 in Khwe) suggest that vivax exerted strong selective pressures of comparable intensity in Madagascar and Africa (Pierron et al. 2018; Kasianov et al. 2025).

## ACKNOWLEDGMENTS

We acknowledge the essential help of the HPC Core Facility of Institut Pasteur for this work. This work was supported by the *Institut Pasteur*, the *Collège de France*, the *Centre Nationale de la Recherche Scientifique* (CNRS), the *Agence Nationale de la Recherche* (ANR) grants LIFECHANGE (ANR 17 CE12 0018 02), CNSVIRGEN (ANR-19-CE15-0009-02) and MORTUI (ANR-19-CE35-0005), the French Government’s *Investissement d’Avenir* program, *Laboratoires d’Excellence* “Integrative Biology of Emerging Infectious Diseases” (ANR-10- LABX-62-IBEID) and “*Milieu Intérieur*” (ANR-10-LABX-69-01), the *Fondation Allianz-Institut de France*, and the *Fondation de France* (no. 00106080).

## DATA ACCESSIBILITY

Supplemental material, including Figures S1 to S14, Tables S1 to S3 (Tables S1-S3 are supplied as a single merged Excel file) can be found with this article online. The data that support the findings of this study are openly available at https://www.simonsfoundation.org/simons-genome-diversity-project/ for the Simons Genome Diversity Project data, at the 1000 Genomes Project web site https://www.internationalgenome.org/ for European and East Asian individuals. The code for simulating our composite scenario, implementing the GAN and the CNN tested in this study will be available at the GitHub code depository site soon (https://github.com/h-e-g/). Because the topology of the CNNs may have various impacts on the estimations, depending on the demographic scenario assumed, we also provide user friendly tools to automatically set the CNN architecture (numbers of convolutions and maxpooling layers, parameters of the convolution, numbers of hidden layers and number of neurons per layer).

## AUTHOR CONTRIBUTIONS

J.C., L.Q.-M and G.L. conceived and designed the study. A.D. and G.L. implemented and validated the CNN. G.L. implemented and validated the GAN. O.P generated the sequence data. G.L. was the lead analyst G.L. with important contributions from A.D., O.P, E.P and J.C. G.L. wrote the manuscript with important contributions from E.P., J.C. and L.Q.-M.

