## Supplementary Figures S1-14 for "Deep learning analyses of DNA sequences resolve the retention of the Duffy-null resistance to Plasmodium vivax malaria in Africa"

**Figure S1. Neutrality statistics.**

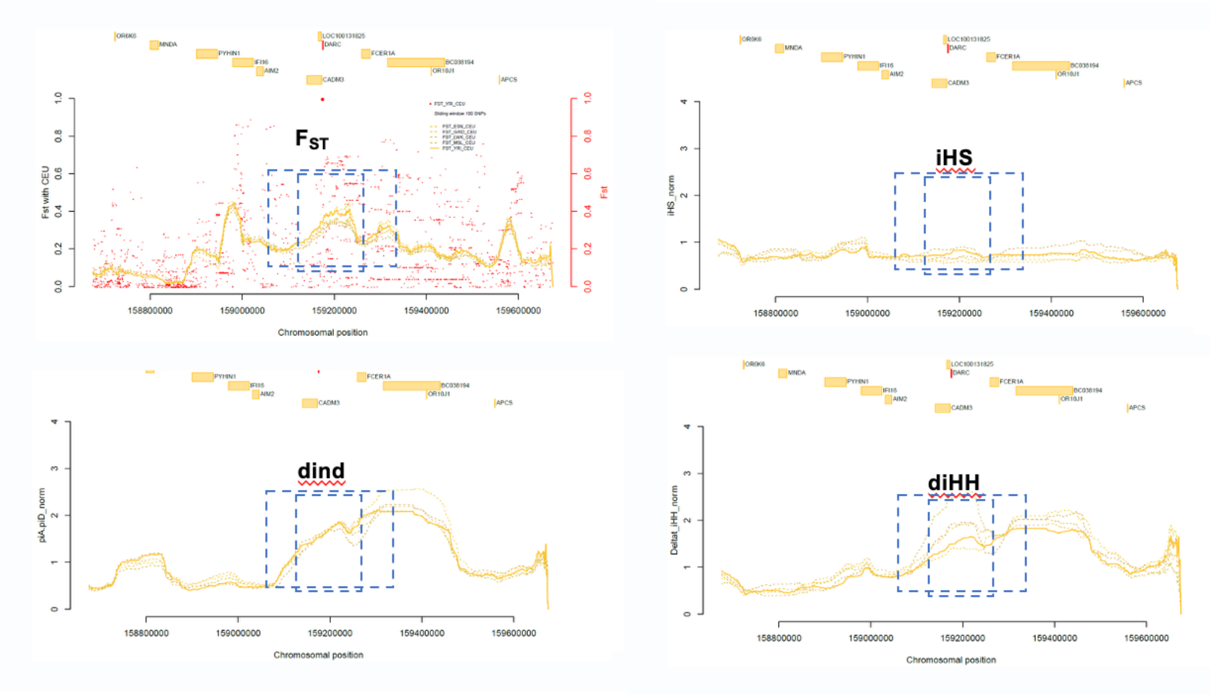


Pattern of the neutrality statistics computed 1Mb around the Duffy-null mutation in the Yoruba population. Dashed blues rectangles display the 100kb and 200kb regions around the Duffy-null variant (rs2814778), in which the proportions of candidate SNPs were computed for each neutrality statistic used in ABC.

**Figure S2. CNN architecture.**

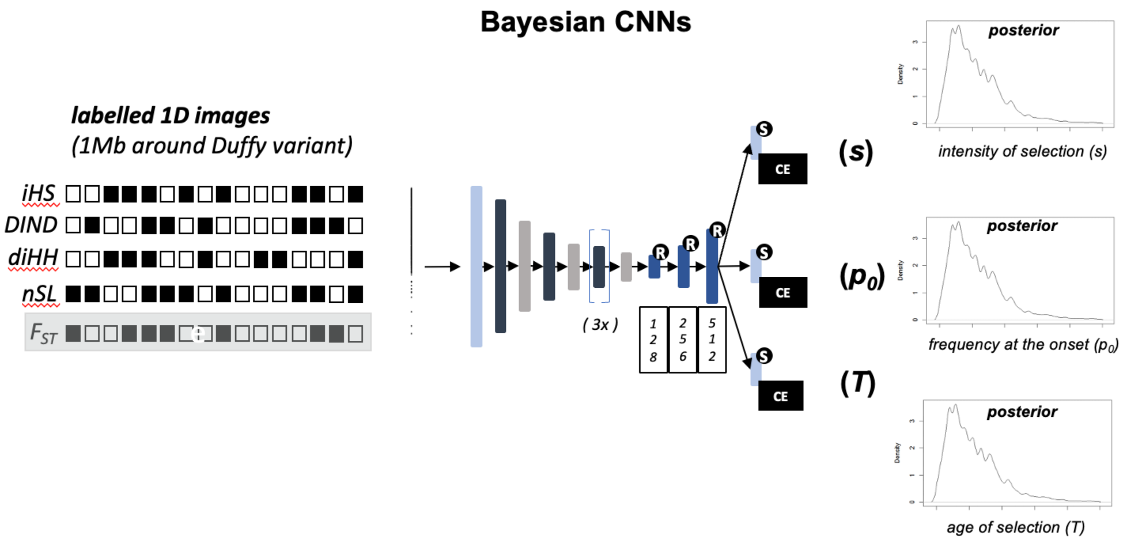


Convolutional neural networks used in this study. Convolution layers with a kernel size of 20 pixels (black), the first convolution layer is connected to the input image, three fully connected layers (bleu, number of neurons are indicated), three output layers (100 neurons each). This CNN was trained with three training epochs.

**Figure S3. GAN architecture.**

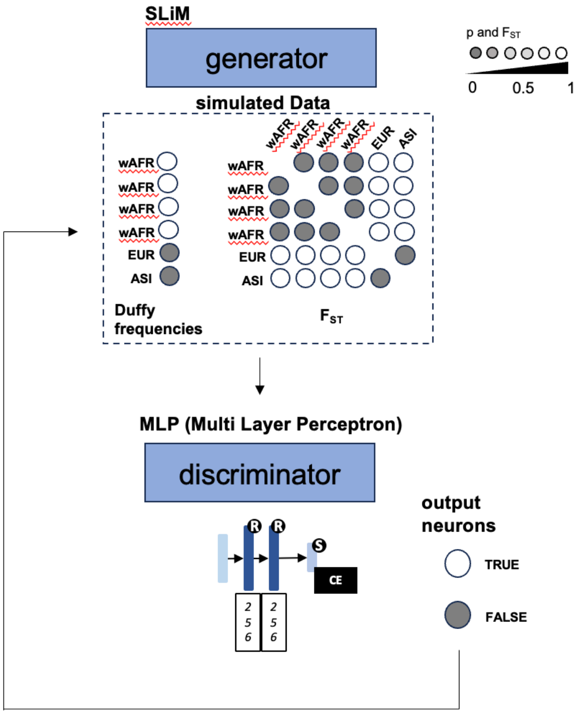


The GAN generator is the SLiM software. Note that here wAFR stand for every African lineage currently living in the western part of Africa (wAFR, wBSP). Multi layer perceptron (MLP) used as discriminator in this study. Two fully connected layers (bleu, number of neurons are indicated), one output layers (2 neurons). This MLP was trained with three training epochs.

**Figure S4. Out of Africa dispersal from the ancestor of western and eastern AFricans.**

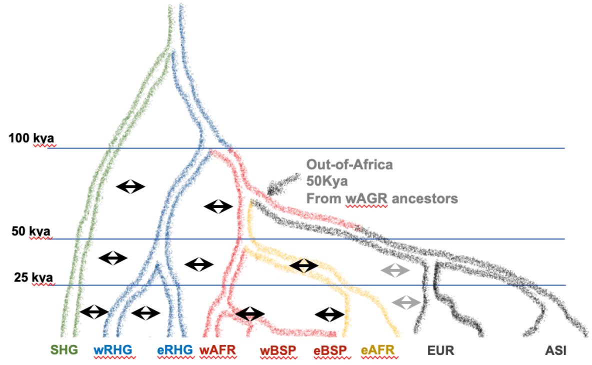


The model close to the Gravel's model used in the study of McManus and colleagues. Same parameter as in the model displayed in Figure 1 (parameter values are given in Table S2). The only difference is an Out of Africa dispersal 50kya but originating from the common ancestor of Bantus speakers, western and eastern Africans (in this case the split between western and eastern Africans was set to 45kya).

**Figure S5. The ancient sweep model.**

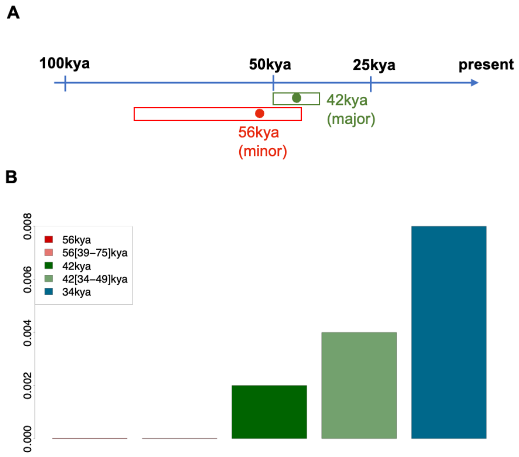


Computer simulations performed using the model close to the Gravel's model displayed in Figure S4. (A) Time ranges of the *T* estimates provided by McManus and colleagues. (B) Percents of simulations declared as "true data" by our trained GAN-discriminator, i.e. simulated data reproducing the Duffy-null fixed in western Africa and absent outside Africa.

**Figure S6. ABC and CNN cross validations.**

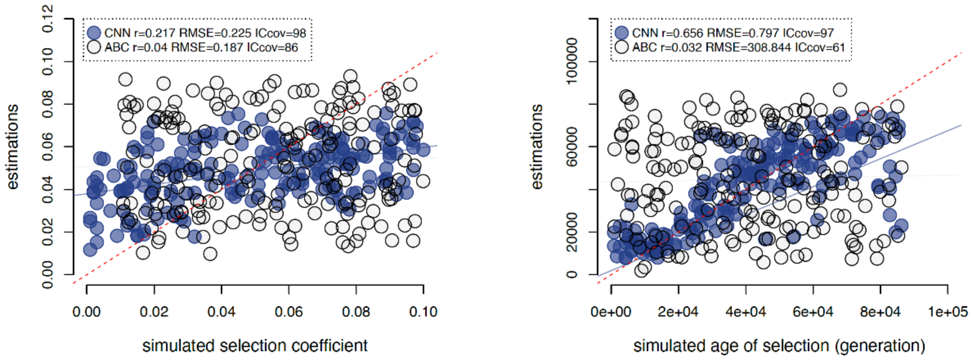


Simulation-based cross validations for the selection coefficient (left panel) and the age of selection (right panel) performed based on 200 pseudo-empirical dataset. Simulations performed assuming frequency at the onset of selection of the order of 0.001, which is in fine the values of $p_0$ estimated for the Duffy-null allele. (Top) Comparisons between the ABC and CNN point estimates with the trues values. The ABC and CNN predictions were performed from 1640 ancient (pseudo-haploids) and 1000 modern (diploids) simulated individuals. Solid colored lines represent the regression lines between true and predicted values obtained for each method. Red dashed solid lines represent the identity line. The linear correlation coefficient *r*, the relative root of the mean square error *RMSE* and the proportion of true values within the credible intervals of estimates *CIcov*, are indicated on each panel.

**Figure S7. GAN-discriminator cross validations.**

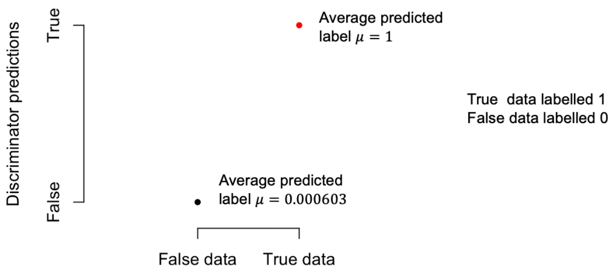


Simulation-based cross validations of the trained GAN-discriminator performed from 10,000 simulated datasets with known labels. Here the label 1 "true data" does not stands for real data, but here the label 1 stands for simulated data with Duffy-null fixed in western Africans and absent in European and Asians, 0 stands for "fake data", i.e. all other cases. The average of predicted label equal 1 showing that our trained GAN-discriminator never failed to recognize "true data". The average of false label equal lower than $10^{-3}$ gives the error rate in this case. Our trained GAN-discriminator may predict false data as being "true data", when the simulated Duffy-null is higher than 0.98 in western Africans and absent in European and Asians.

**Figure S8. Posterior distributions of the intensity of selection *s*.**

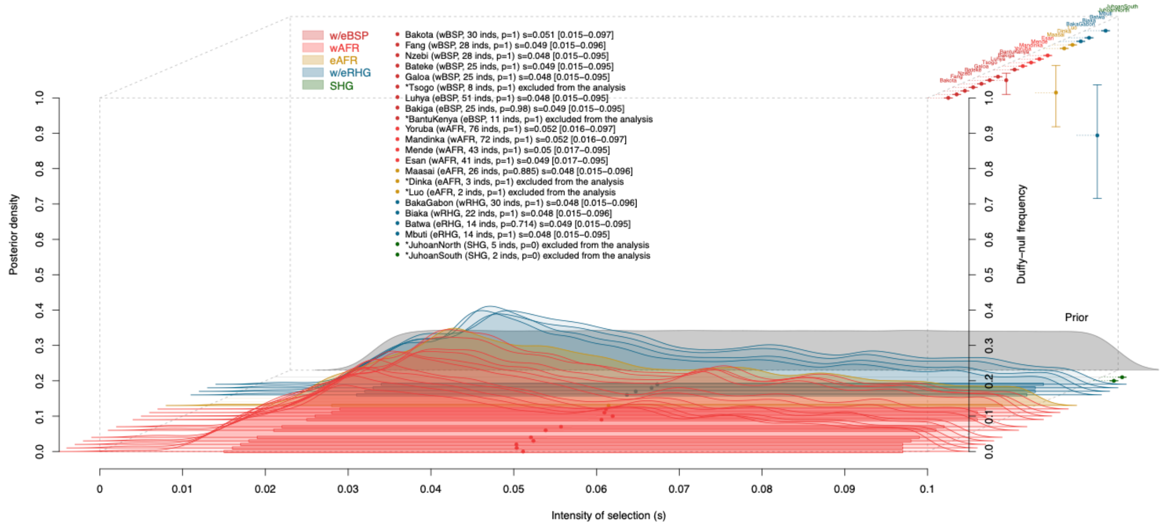


Same legend as in Figure 3A.

**Figure S9. Posterior distributions of the frequency at the onset of selection $p_0$.**

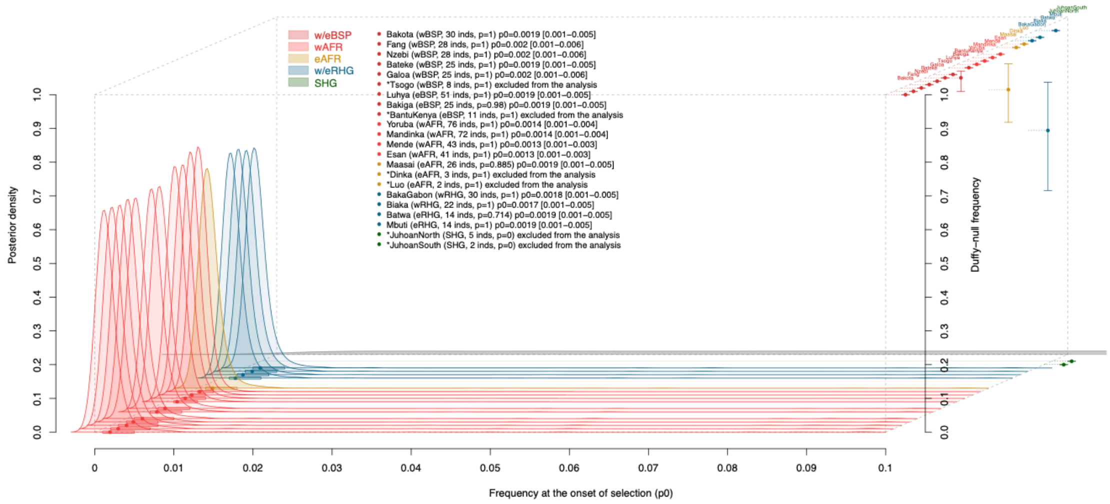


Same legend as in Figure 3A.

**Figure S10. Haplotypes in the eastern agro-pastoralist population (eAFR) Maasai.**

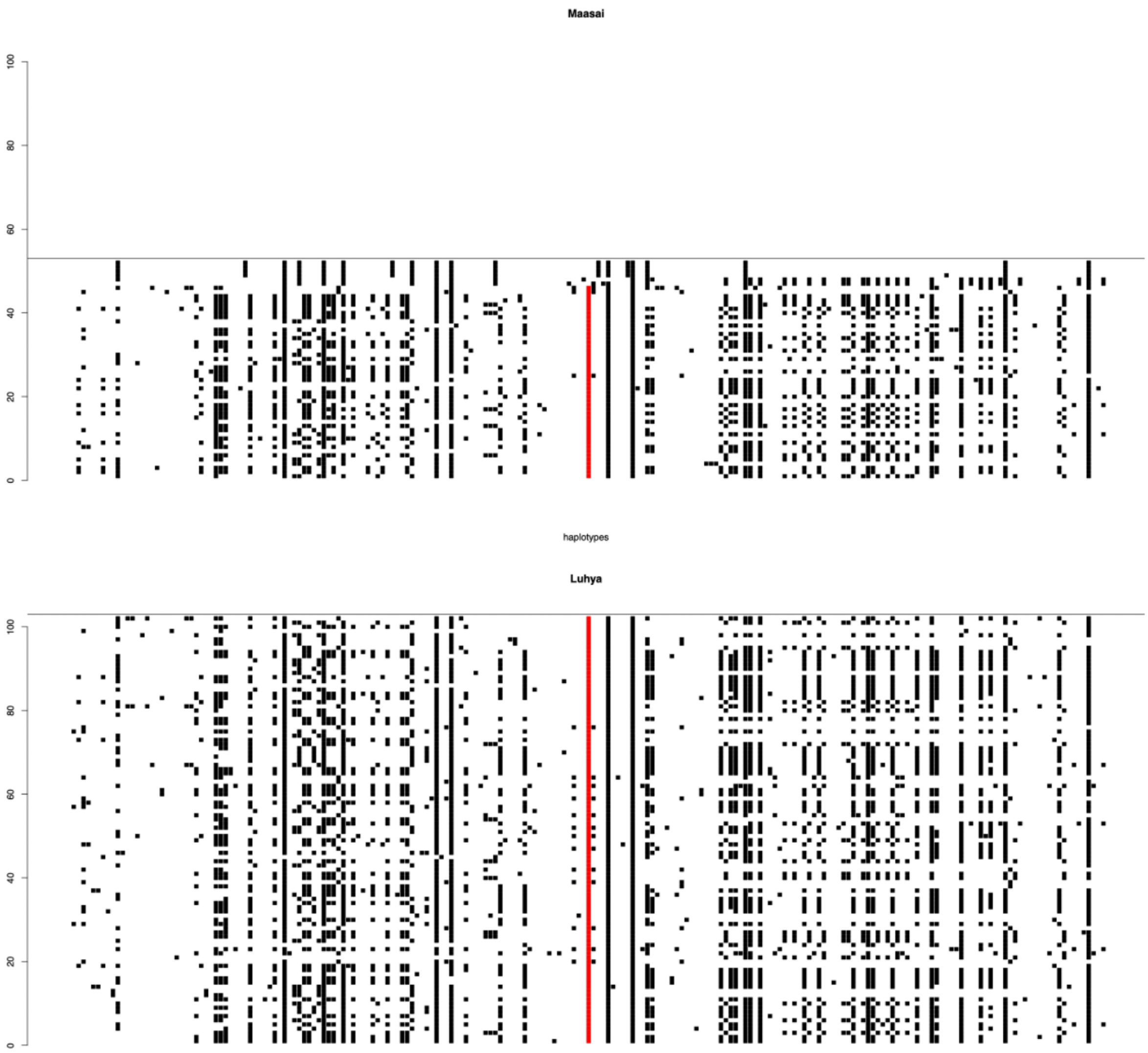


(Top) Haplotypes 40Kb around the Duffy-null allele displayed in red. (Bottom) Haplotypes observed in the putative parental Bantu speaking population used as reference are also displayed.

**Figure S11. Haplotypes in the eastern hunter-gatherer population (eRHG) Batwa.**

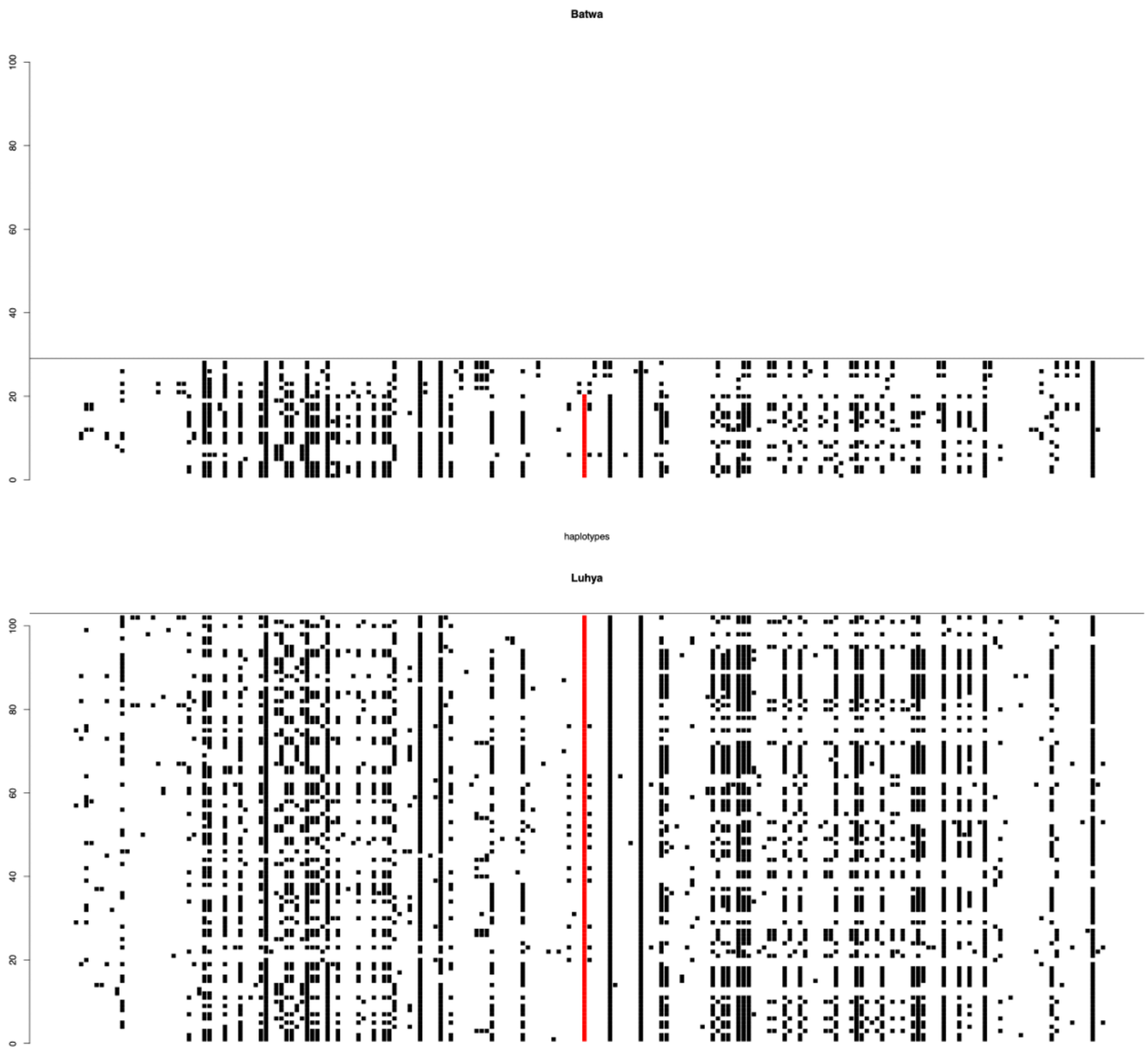


(Top) Haplotypes 40Kb around the Duffy-null allele displayed in red. (Bottom) Haplotypes observed in the putative parental Bantu speaking population used as reference are also displayed.

**Figure S12. Haplotypes in the eastern hunter-gatherer population (eRHG) Mbuti.**

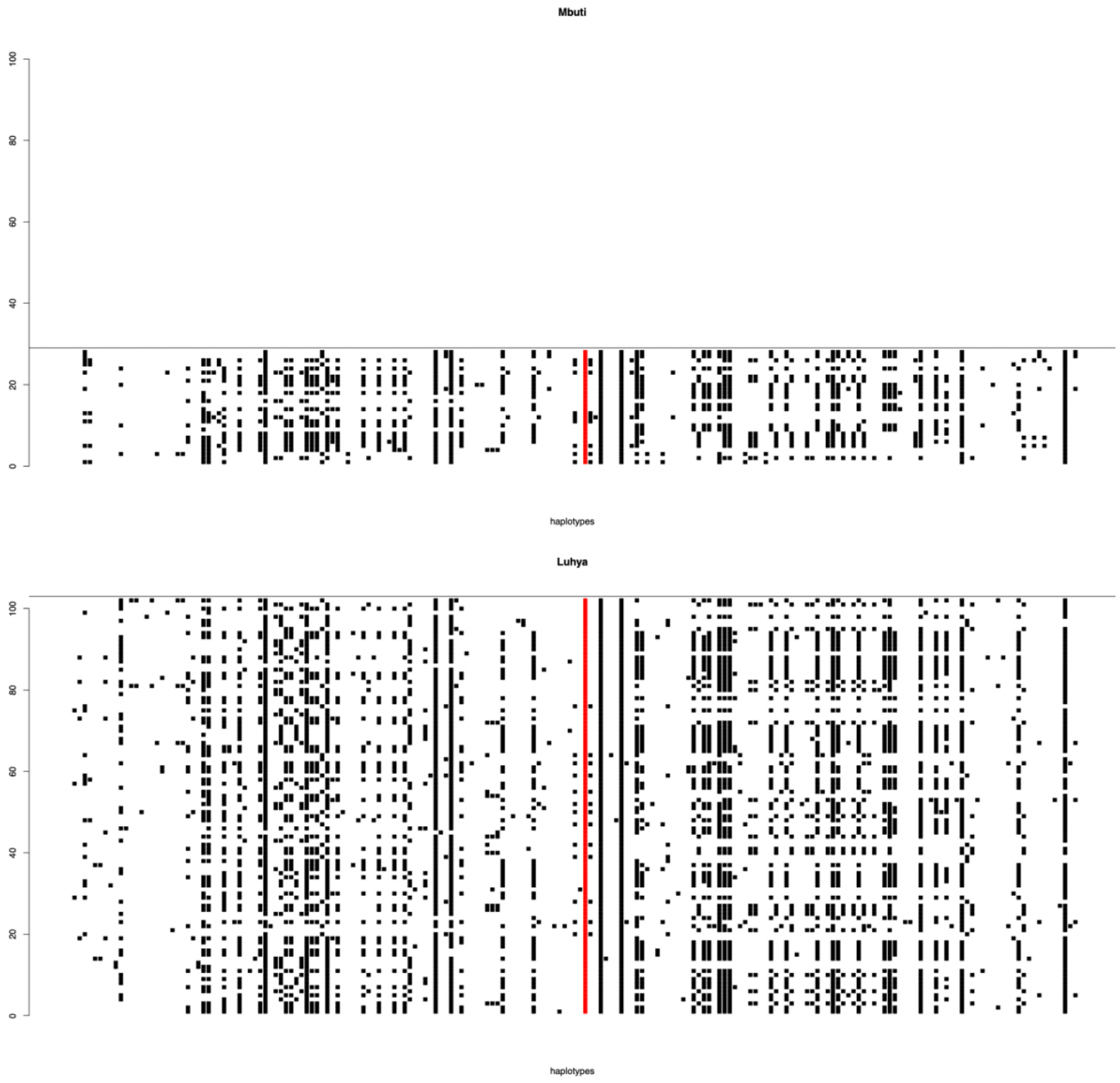


(Top) Haplotypes 40Kb around the Duffy-null allele displayed in red. (Bottom) Haplotypes observed in the putative parental Bantu speaking population used as reference are also displayed.

**Figure S13. Haplotypes in the western hunter-gatherer population (wRHG) BakaGabon.**

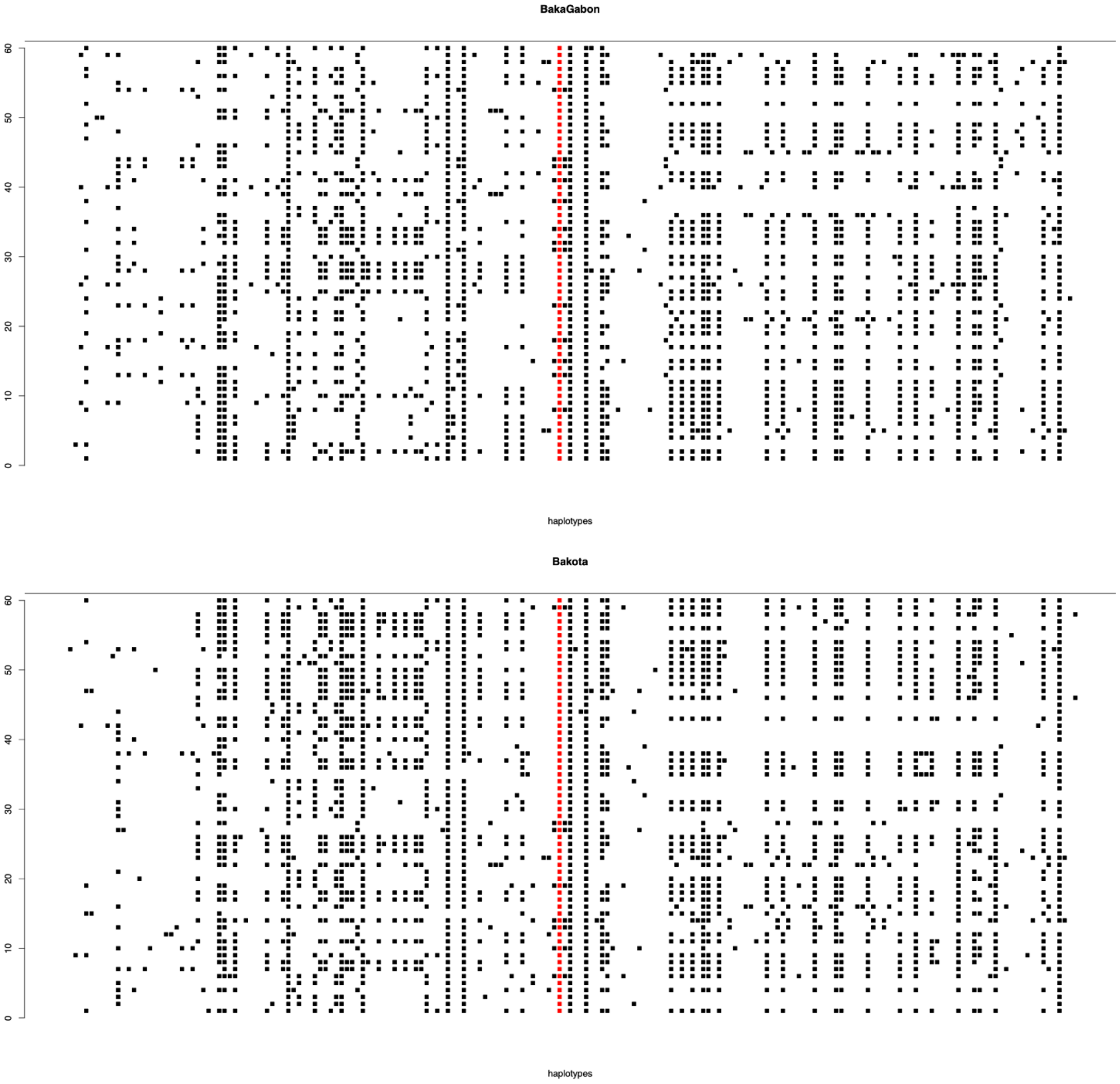


(Top) Haplotypes 40Kb around the Duffy-null allele displayed in red. (Bottom) Haplotypes observed in the putative parental Bantu speaking population used as reference are also displayed.

**Figure S14. Haplotypes in the western hunter-gatherer population (wRHG) Biaka.**

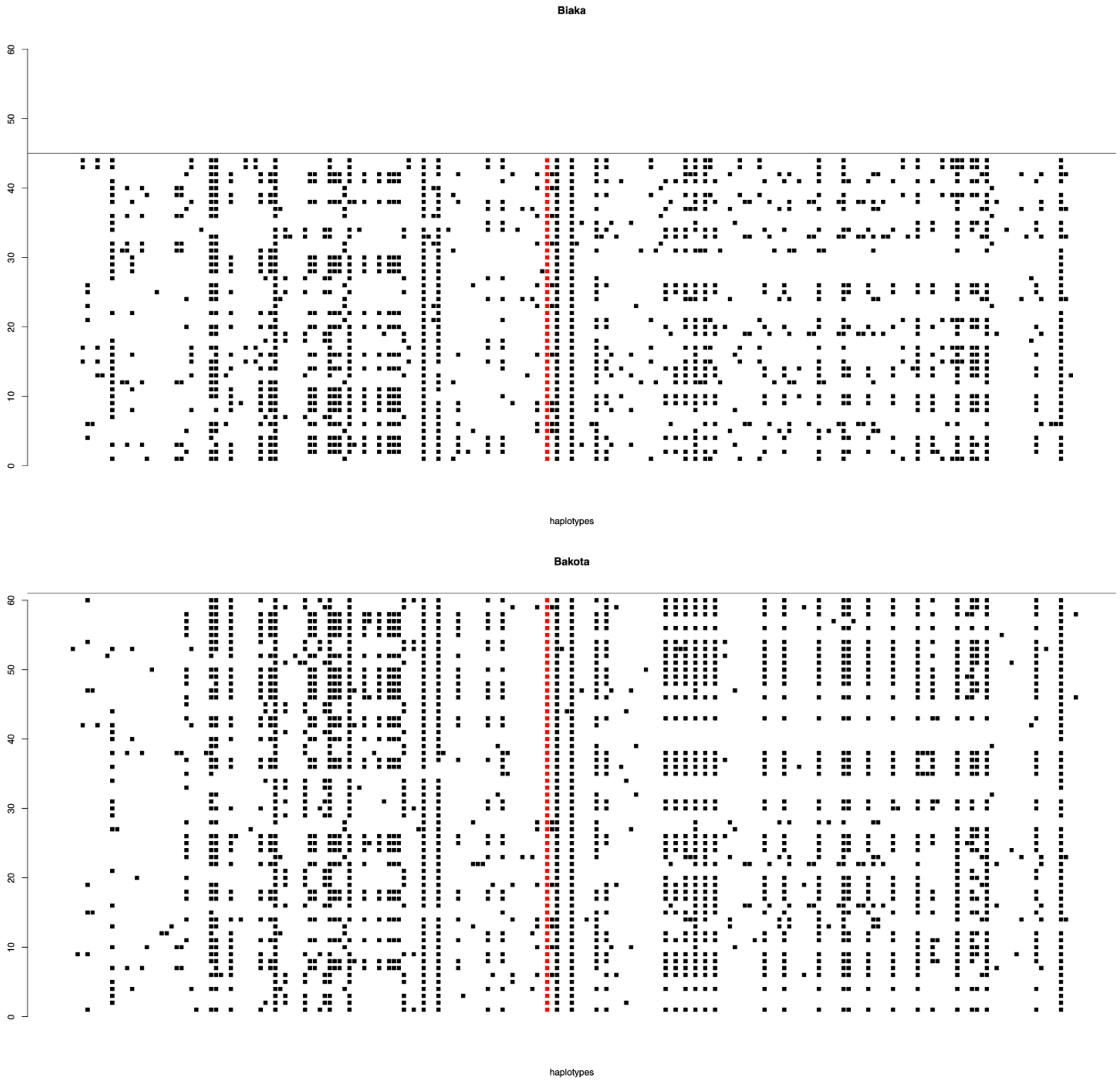


(Top) Haplotypes 40Kb around the Duffy-null allele displayed in red. (Bottom) Haplotypes observed in the putative parental Bantu speaking population used as reference are also displayed.

**Table S1. List of analyzed populations**

**Table S2. Parameters of the simulated models**

**Table S3. Number of differences between haplotypes**

Tables S1-3 are supplied as Excel files
